# Within-host evolutionary dynamics and tissue compartmentalization during acute SARS-CoV-2 infection

**DOI:** 10.1101/2022.06.21.497047

**Authors:** Mireille Farjo, Katia Koelle, Michael A. Martin, Laura L. Gibson, Kimberly K.O. Walden, Gloria Rendon, Christopher J. Fields, Fadi G. Alnaji, Nicholas Gallagher, Chun Huai Luo, Heba H. Mostafa, Yukari C. Manabe, Andrew Pekosz, Rebecca L. Smith, David D. McManus, Christopher B. Brooke

## Abstract

The global evolution of SARS-CoV-2 depends in part upon the evolutionary dynamics within individual hosts with varying immune histories. To characterize the within-host evolution of acute SARS-CoV-2 infection, we deep sequenced saliva and nasal samples collected daily from immune and unvaccinated individuals early during infection. We show that longitudinal sampling facilitates high-confidence genetic variant detection and reveals evolutionary dynamics missed by less-frequent sampling strategies. Within-host dynamics in both naïve and immune individuals appeared largely stochastic; however, we identified clear mutational hotspots within the viral genome, consistent with selection and differing between naïve and immune individuals. In rare cases, minor genetic variants emerged to frequencies sufficient for forward transmission. Finally, we detected significant genetic compartmentalization of virus between saliva and nasal swab sample sites in many individuals. Altogether, these data provide a high-resolution profile of within-host SARS-CoV-2 evolutionary dynamics.

## Introduction

The large-scale sequencing and phylogenetic analyses of clinical samples during the SARS-CoV-2 pandemic have captured global evolutionary dynamics of the virus with unprecedented speed and resolution. However, our understanding of viral evolutionary dynamics within individual infected hosts remains limited. Most studies of SARS-CoV-2 within-host evolution have focused on chronic infections of immunocompromised individuals, as these patients are more amenable to repeated, longitudinal sampling. It has been hypothesized that chronic infections promote the emergence of novel viral variants by providing a combination of prolonged time for replication and relatively weak immune selection that promotes the emergence of variants with increased fitness to high frequency within the host (Avanzato et al., 2020; Baang et al., 2021; Corey et al., 2021). Persistent replication within immunocompromised individuals treated with convalescent sera or therapeutic monoclonal antibodies has also been identified as a potential source of antigenically novel variants (Choi et al., 2020; Kemp et al., 2021; Truong et al., 2021).

Previous studies of SARS-CoV-2 within-host evolutionary dynamics during acute infection of immunocompetent hosts observed low within-host diversity in SARS-CoV-2 populations, with most specimens containing 15 or fewer intra-host single-nucleotide variants (iSNVs) (Braun et al., 2021; Lythgoe et al., 2021; Tonkin-Hill et al., 2021; Valesano et al., 2021). Studies of household transmission reaffirm that within-host diversity is low and that iSNVs are rarely transmitted between members of a household (Braun et al., 2021; Lythgoe et al., 2021; Valesano et al., 2021). Altogether, these data suggest that acute infections typically exhibit low overall levels of within-host genetic diversity and that the selection-driven emergence of iSNVs to high frequency during acute infection is likely rare. However, our understanding of within-host evolutionary dynamics has been hampered by the absence of high-resolution time course data within individuals.

The extent to which pre-existing immunity, elicited either through vaccination and/or prior infection, influences the within-host evolution of SARS-CoV-2 is poorly understood. For two-dose vaccinations, it remains unclear whether administration of a single dose without a follow-up may create an evolutionary sandbox where moderate immune selection in the absence of rapid clearance can drive the emergence of immune-escape variants (Cobey et al., 2021; Saad-Roy et al., 2021). A similar question has been raised by the emergence of new variants like Omicron that are able to efficiently replicate in vaccinated individuals where the virus may accumulate additional immune escape substitutions. Thus, it is important to characterize the extent of immune selection and potential for escape variant emergence during infections of immune-competent individuals at differing stages of vaccination.

To characterize viral evolutionary dynamics during acute SARS-CoV-2 infection, we sequenced longitudinal nasal swab and saliva samples collected from 32 students, faculty, and staff at the University of Illinois at Urbana-Champaign enrolled during the early stages of infection through an on-campus screening program (Ranoa et al., 2021). This cohort included 20 naive individuals and 12 individuals with presumed pre-existing immunity to SARS-CoV-2 resulting from vaccination or prior infection. By taking repeated measures of iSNV frequencies from two sample sites (mid-turbinate (MT) nasal swab and saliva) within individuals, we were able to generate high-resolution profiles of iSNV dynamics between tissue compartments and across time. Our results demonstrate that selection, genetic drift, and spatial compartmentalization all play important roles in shaping the within-host evolution of SARS-CoV-2 populations.

## Results

### Sample collection

During the 2020-2021 school year, all students, faculty, and staff on the University of Illinois at Urbana-Champaign campus were required to undergo saliva-based PCR testing for SARS-CoV-2 at least twice a week (Ranoa et al., 2021). We enrolled individuals who were either (a) within 24 hours of their first positive test result, or (b) within 5 days of exposure to someone else who tested positive. Daily saliva samples and nasal swabs were collected from each enrolled participant for up to 14 days. Details on the dynamics of viral shedding in this cohort have been published previously (Ke et al., 2022a, 2022b; Smith et al., 2021).

### Optimization and validation of saliva sample sequencing protocol

The saliva-based PCR assay used in this study involves a 30-minute treatment at 95°C which partially degrades the viral RNA present in the sample and could potentially compromise sequencing quality (Ranoa et al., 2020, 2021). To address this concern and determine whether saliva Ct values are predictive of sequencing data quality, we examined sequencing depth across samples with a range of Ct values. Over a set of 10 samples that spanned a Ct range of 21.63 to 34.34, we observed a clear negative correlation between N gene Ct value and coverage depth (Adjusted R-squared = 0.4296, p = 0.02359) (**Fig 1A**). For Ct values below 28, we obtained average per-nucleotide read depths of over >10,000 reads, indicating that high quality sequence data can be obtained from heat-treated samples.

**Figure 1:**
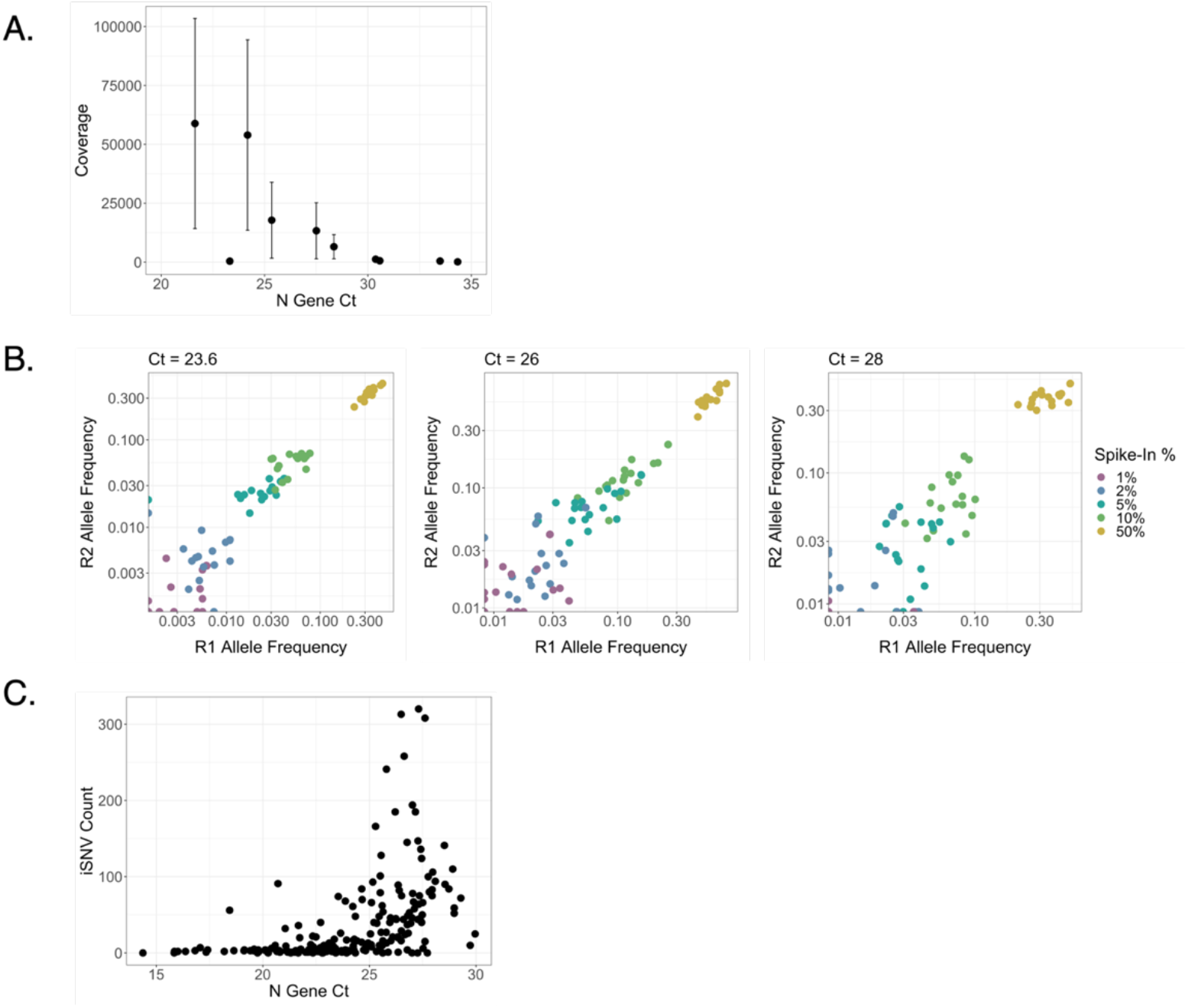
Relationship between saliva sample Ct values and sequence quality. **(A)** Linear regression between Ct values of nucleocapsid (N) gene and mean sequence coverage depth. Error bars represent standard deviation. **(B)** Frequencies of characteristic B. 1.1.7 SNPs at Ct values of 23.6, 26, and 28. B. 1.1.7 RNA was spiked into B. 1.2 RNA at final percentages ranging from 1% to 50% and divided between two replicates (R1 and R2). **(C)** Relationship between Ct of N and total iSNV count in resulting sequences (Spearman’s rank correlation, rho = 0.6351, p < 0.001).

We next evaluated the relationship between Ct values and the reliability of iSNV detection in saliva samples. We generated control samples in which RNA isolated from a B.1.1.7 (Alpha) lineage sample was spiked into a B.1.2 lineage sample at defined frequencies of 50%, 10%, 5%, 2%, and 1%. We normalized both B.1.1.7 and B.1.2. samples to Ct values of 23.6, 26, or 28 based on Ct values prior to mixing. Spike-ins were then divided into replicate samples and deep-sequenced. We compared the measured frequencies of the 17 characteristic B.1.1.7 SNPs and indels between technical replicates (**Fig 1B**). We detected B.1.1.7-associated mutations at the expected frequencies for spike-ins at 5% or greater, but frequency estimates were much noisier for the 1% and 2% spike-ins. The correlation between technical replicates was also stronger at dilutions above 2% and in samples with a Ct of 23.59 or 26 than in samples with a Ct of 28. Based off these results, we set a variant calling threshold of 3% and a Ct cutoff of 28 for analyzing saliva samples.

We ranked study participants based on the number of saliva samples with Ct<28 and the range of time that these samples covered and selected 20 unvaccinated participants for further study. We also selected 12 study participants who were either vaccinated (fully or partially; definitions in methods) or reported a previous positive SARS-CoV-2 PCR test result and had at least one saliva sample with Ct<30. We refer throughout to this group as “immune” as we assume they mounted some sort of adaptive immune response to vaccination or infection; however, we were unable to empirically measure immune responses in this study. We chose a higher Ct threshold for these immune participants because Ct values overall were much higher in this group (Ke et al., 2022a, 2022b). Across the entire cohort, the number of iSNVs per sample was correlated with the Ct value of the sample (Spearman’s rank correlation, rho = 0.6351, p < 0.001), further demonstrating that high Ct values can contribute to noise in iSNV detection (**Fig 1C**).

### Analysis of within-host diversity

We next examined the diversity within and between individual saliva samples, focusing on iSNVs and short insertions/deletions (indels) present at frequencies between 3-97%, with coverage depths of >1000 reads. The numbers of iSNVs that fit these criteria varied substantially between samples, generally spanning values between 1 and 100 at different points during infection (**Fig 2A,B**). To minimize false positive iSNV calls, we focused on iSNVs that appeared in at least two saliva samples collected from a given individual across different dates of infection (shared iSNVs). Numbers of shared iSNVs were similar between participants, averaging 6.31 shared iSNVs per individual, which aligns with previous assessments (Braun et al., 2021; Tonkin-Hill et al., 2021; Valesano et al., 2021).

**Figure 2:**
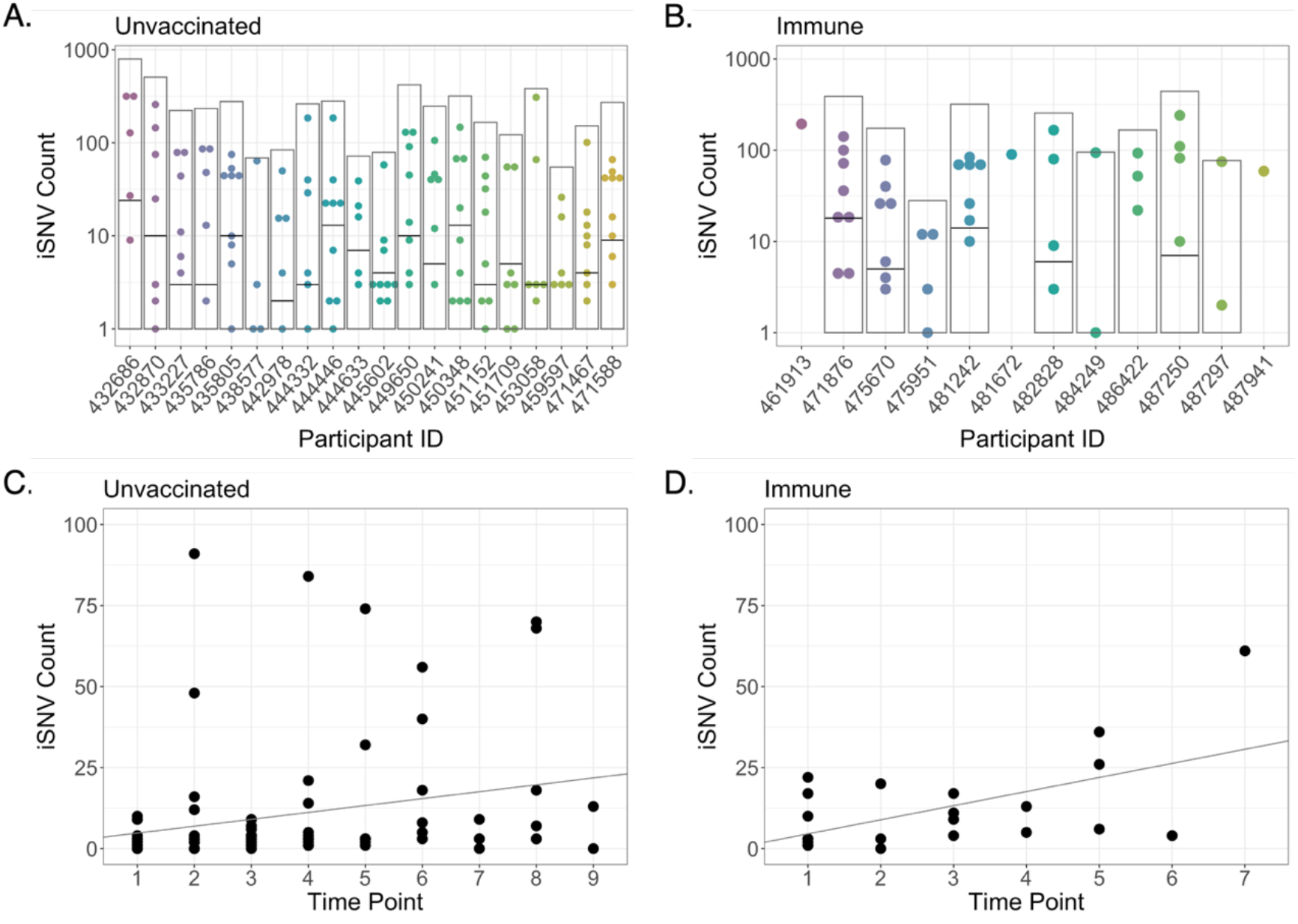
Intra-host single nucleotide variant (iSNV) diversity compared between samples and individuals. **(A)** Total iSNV counts for each sample from each unvaccinated participant. Light grey boxes indicate total iSNV count for all samples and horizontal black lines indicate number of shared iSNVs for each participant. **(B)** iSNV counts for immune participants. **(C)** iSNV counts for individual samples with Ct < 25 from naïve participants as a function of number of days post-enrollment (Adjusted R-squared = 0.05007, p = 0.02255). Line represents linear regression. **(D)** iSNV counts for individual samples with Ct < 25 from immune participants as a function of number of days post-enrollment (Adjusted R-squared = 0.2857, p = 0.006359). Line represents linear regression.

Naive participants had lower overall iSNV counts than immune participants, with average variant counts of 33.39 and 51.73 respectively (Welch two sample t-test, p = 0.05645) (**Fig 2A,B**). However, the higher iSNV counts in immune samples is likely due to higher Ct values, as indicated above (unvaccinated average = 23.79, immune average = 25.15, p = 0.007773). There was no significant difference in the number of shared iSNVs between the two groups (unvaccinated average = 6.65, immune average = 5.56, Welch two sample t-test, p = 0.6734). We also observed an upward trend in iSNV counts over time in both groups. To account for the impact of high Ct values on artifactual iSNV accumulation, we only considered samples with a Ct below 25 in our analysis (we did not restrict the analysis to only shared iSNVs, to avoid violating assumptions of independence between data points). We found a stronger correlation between iSNV counts and time of sample collection in immune individuals (Adjusted R-squared = 0.2857, p = 0.006359) than in naïve individuals (Adjusted R-squared = 0.05007, p = 0.02255), but the relationship was significant in both groups (**Fig 2C,D**). These data indicate similar overall levels of within-host diversity in naïve and immune individuals, but potentially a higher rate of mutant accumulation over time in immune individuals.

We next examined the distributions of shared iSNVs across the viral genome for both unvaccinated and immune populations. Since 3 immune individuals only had a single timepoint, and 4 others had no shared iSNVs, we excluded these 7 immune participants from this analysis. We also detected four frameshift mutations (at nucleotide positions 6696, 11074, 15965, and 29051) in many samples at low but consistent frequencies. Given the low likelihood of identical frameshift mutations repeatedly arising and persisting in multiple populations, we concluded that these variants are likely sequencing artifacts and we removed them from the dataset.

After the removal of these variants, we still observed several iSNVs and indels that recurred across multiple naïve individuals, including a t29760c substitution in the 3’ UTR region present in 9/20 naive participants, and several coding substitutions in ORF1ab (**Fig 3A**). In immune individuals, a handful of mutations were shared by pairs of participants (**Fig 3B**). These included a P681H substitution at the S1/S2 cleavage site in the spike protein associated with the B.1.1.7 (Alpha) lineage and mutations (a G➔A substitution and a G➔GAACA insertion) at nucleotide position 28262, in the untranslated region between the E (Envelope) and N (Nucleocapsid) genes. From our data, we cannot determine whether shared mutations arose independently in multiple participants or were transmitted. Beyond these exceptions, the vast majority of iSNVs were only detected in a single study participant (**Fig S1**).

**Figure 3:**
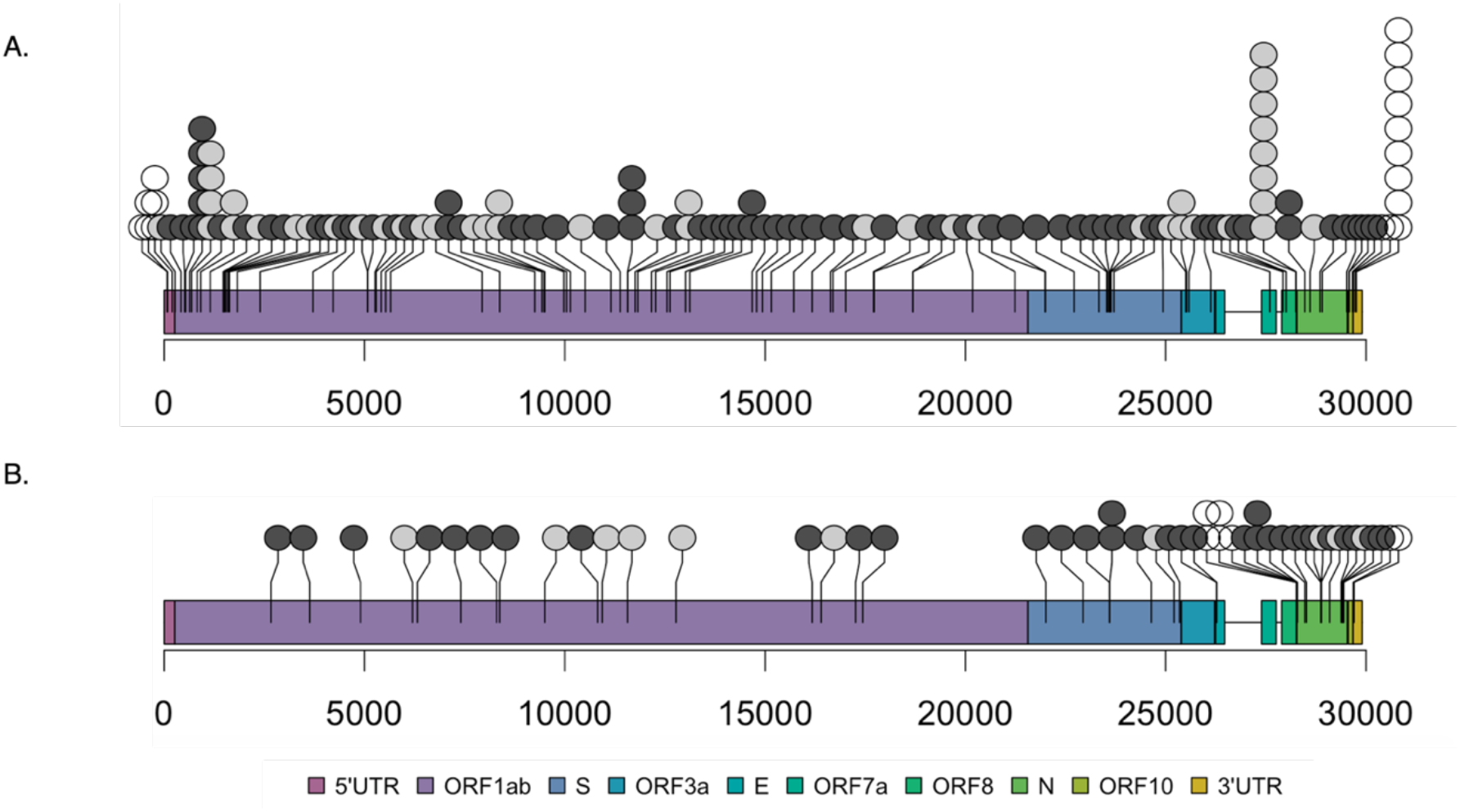
Locations of shared iSNVs across the SARS-CoV-2 genome. Genome locations of shared iSNVs found in naive **(A)** and immune **(B)** participants. Number of dots at a locus indicate number of participants in which the shared iSNV was detected. Light grey dots indicate synonymous mutations, dark grey dots indicate nonsynonymous mutations, and white dots indicate UTR mutations.

Mutations were not evenly distributed across the viral genome and clear hotspots of accumulation were easily observable. In naive participants, we observed 6 mutations between ORF1ab positions 402-457 and 4 mutations between spike positions 655-681, near the S1/S2 cleavage site (**Fig 3A**). As indicated above, S:P681H was also observed in two members of the immune cohort. In immune individuals, we observed 3 clusters of mutations in the N gene (positions 3-80, 199-204, and 370-391; **Fig 3B**). Mutations in the N gene were enriched in immune participants – they made up 31.37% of shared variants in immune individuals but only 4.29% of shared variants in naïve participants (Fisher’s exact test, p < 0.001). This enrichment was largely driven by samples from one vaccinated individual, who had 11 shared iSNVs in the N gene. However, when the individual was removed from the study set, immune individuals still exhibited higher proportions of N iSNVs than naïve individuals (Fisher’s exact test, p = 0.03669). This was surprising because the N protein is not targeted by currently licensed vaccines. Despite similarities in overall levels of within-host diversity between the two groups, there appear to be differences between naive and immune individuals in the distribution of this diversity across the viral genome.

### Compartmentalization between tissue environments

Previous studies revealed that SARS-CoV-2 replication dynamics can be highly discordant between saliva and nasal swab samples, suggesting strong compartmentalization of virus between different anatomical sites (Ke et al., 2022a, 2022b). To directly evaluate the extent of compartmentalization between nasal and saliva-associated tissue sites, we compared iSNV frequencies between paired saliva and nasal swab samples over the course of infection in 13 individuals with high quality sequences for both saliva and nasal samples. We first simply compared the frequencies of shared iSNVs present at any frequency in saliva versus nasal swab samples (**Fig 4A**). In the absence of compartmentalization, we would expect iSNV frequencies to be highly correlated between sample sites. Instead, data points almost exclusively fell along the edges of the plot, consistent with substantial compartmentalization between sample sites.

**Figure 4:**
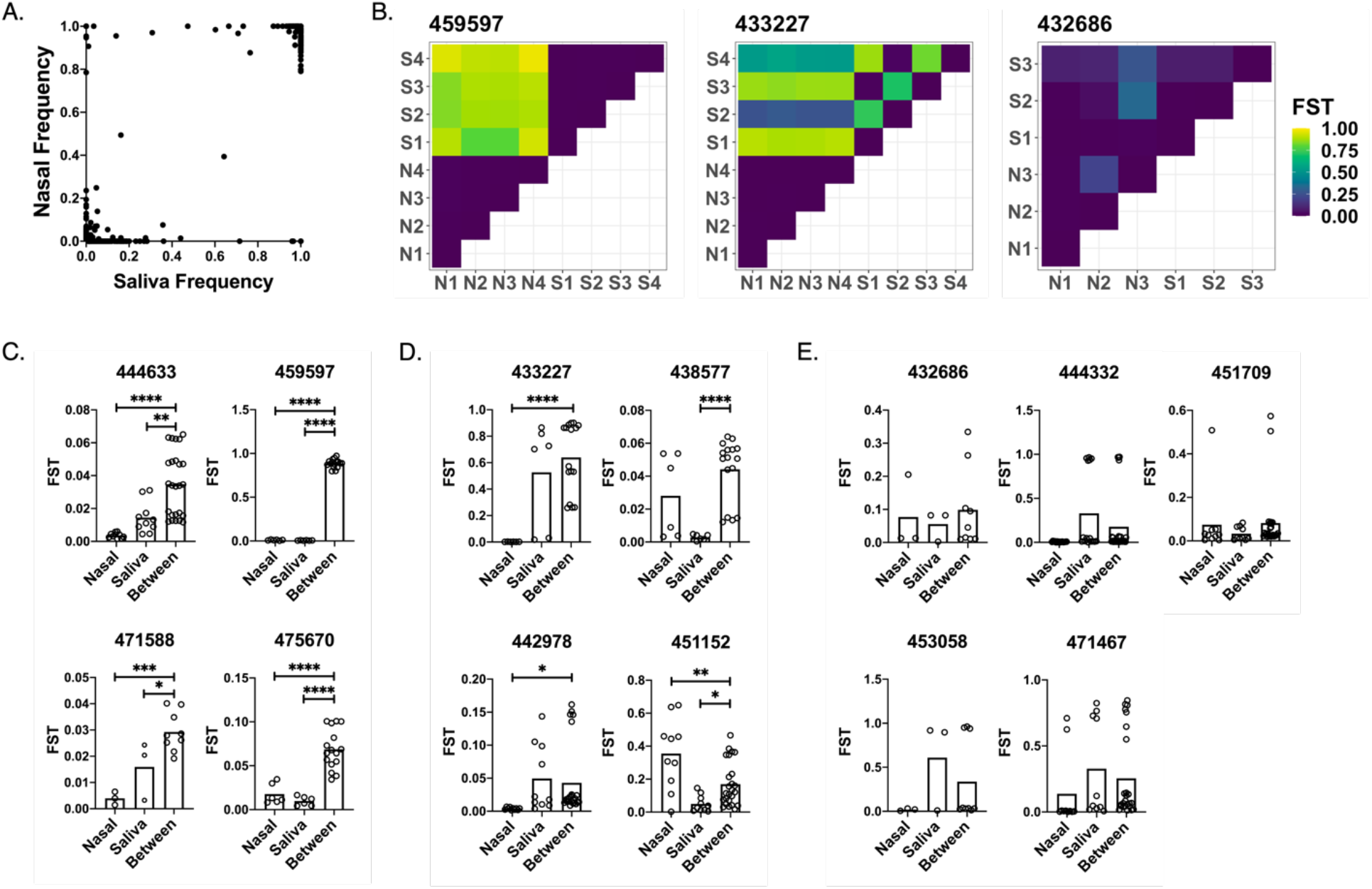
Quantification of genetic compartmentalization of virus between sample sites. **(A)** Comparison of iSNV frequencies between matched samples in nasal and saliva environments. **(B)** Representative heatmaps exemplifying strong (459597), partial (433227), and insignificant (432686) compartmentalization. Maps show F_ST_ values between pairs of samples from nasal (“N”) and/or saliva (“S”) environments (numbered by order of sampling). **(C)** Participants exhibiting strong compartmentalization (within-nasal and within-saliva F_ST_ values are significantly lower than F_ST_ values from paired nasal-saliva samples). **(D)** Participants exhibiting partial compartmentalization (one set of within-environment F_ST_ values is lower than between-environment F_ST_ values). **(E)** Participants exhibiting no significant compartmentalization (neither set of within-environment F_ST_ values is lower than between-environment F_ST_ values). Asterisks indicate levels of significance (* = p < 0.05, ** = p < 0.01, *** = p < 0.001, **** = p < 0.0001). P-values are derived from unpaired t-tests between each group.

To quantify the extent of compartmentalization more precisely, we calculated fixation indices (F_ST_) within and between environments (**Fig 4B,C,D**). The fixation index measures the ratio of allele frequency variation between sub-populations versus the variation in the total population. F_ST_ values range from 0 to 1, and values closer to 1 indicate higher levels of variation between populations. In 7 out of 13 individuals (and overall), the between-environment F_ST_ values were significantly higher than the within-nasal F_ST_ values, reflecting compartmentalization. In 6 out of 13 individuals (but not overall), the between-environment F_ST_ values were significantly higher than the within-saliva F_ST_ values (**Fig 4C,D,E**), again indicative of genetic compartmentalization between tissue compartments.

Study participants fell into three subcategories: (1) higher variation between nasal and saliva environments than within either environment, consistent with strong tissue compartmentalization (**Fig 4B,C**); (2) higher variation between environments than within one environment, consistent with partial compartmentalization (**Fig 4B,D**); and (3) no difference in between-environment versus within-environment variation, consistent with the absence of significant compartmentalization (**Fig 4B,E**). There was only one instance (participant 451152) where within-environment F_ST_ values were significantly higher than between-environment F_ST_ values (**Fig 4**). Our data suggest a significant degree of genetic compartmentalization between tissue environments present in most (8/13), but not all, participants examined.

### Within-host evolutionary dynamics

Our dense longitudinal sampling allowed us to examine the evolutionary forces shaping SARS-CoV-2 populations over the course of acute infection. We first compared numbers of nonsynonymous to synonymous mutations (within a frequency range of 0.03 to 0.97, including both shared and unshared iSNVs) for each saliva sample from each participant (**Fig 5A,B**). We normalized nonsynonymous and synonymous mutation counts based on estimates of total numbers of nucleotide positions across the SARS-CoV-2 genome where a substitution would have a protein-coding effect or a silent effect, respectively. We did not detect any significant temporal trend in dN/dS ratios in either unvaccinated or immune individuals (**Fig 5A,B**). Overall, dN/dS ratios were significantly higher in immune individuals compared with naive individuals, with mean values of 3.411 and 2.664, respectively (Welch two sample t-test, p = 0.02966)(**Fig 5C**).

**Figure 5:**
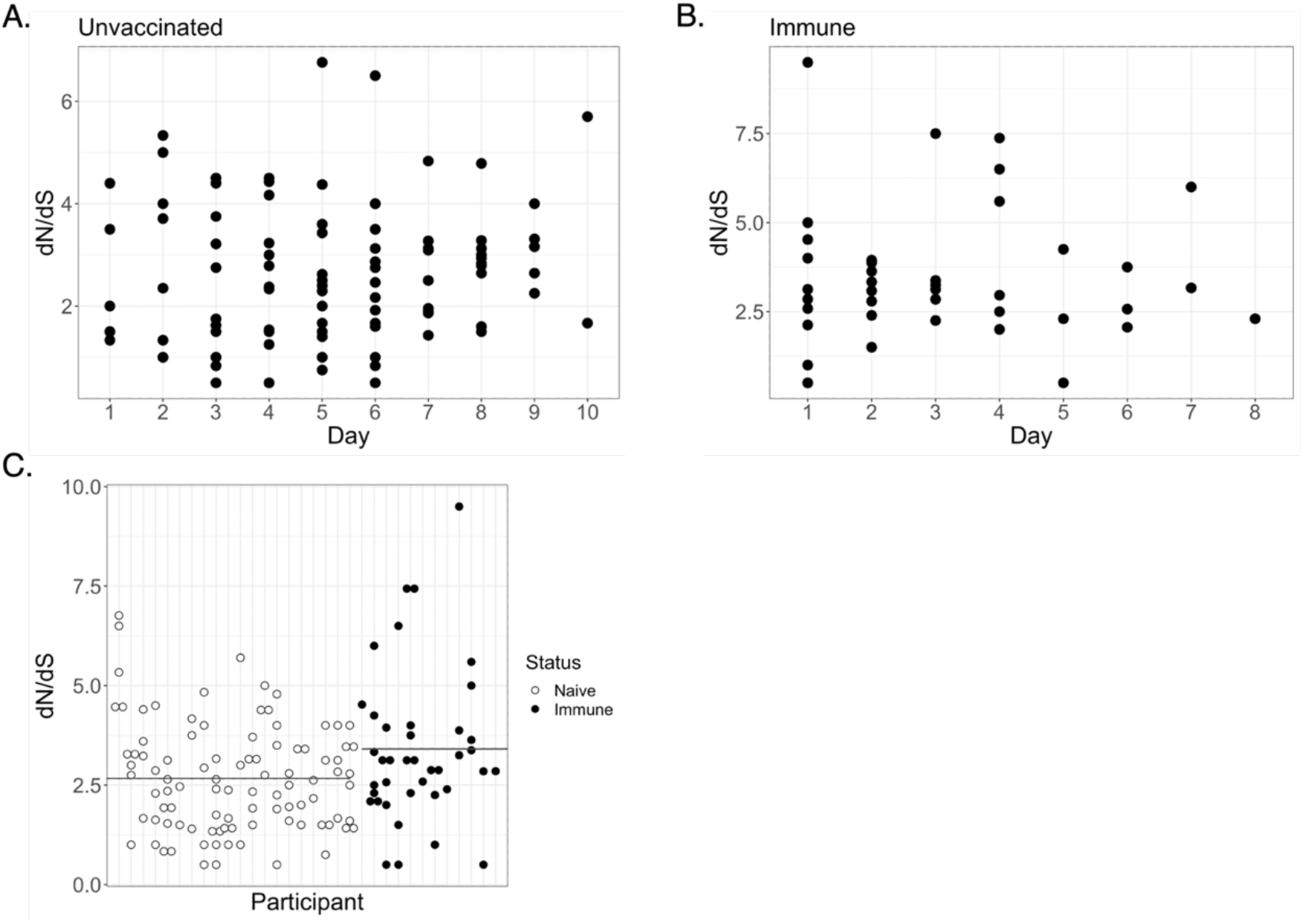
dN/dS ratios significantly differ between naïve and immune individuals. **(A)** dN/dS values for all unvaccinated participants over time. **(B)** dN/dS values for all immune participants over time. **(C)** Comparison between dN/dS values for unvaccinated and immune participants (unvaccinated mean = 2.664, immune mean = 3.411, p = 0.02966).

While there is not a clear relationship between dN/dS ratios obtained from individual related populations and the evolutionary forces acting on those populations (Kryazhimskiy and Plotkin, 2008), our data clearly show an enrichment for nonsynonymous iSNVs within SARS-CoV-2 infected hosts that is more pronounced within immune individuals. This pattern may reflect positive selection occurring within immune individuals; alternatively, it may reflect higher levels of genetic drift in immune individuals (potentially due to lower overall viral loads) and the resulting inability of purifying selection to act as efficiently in these individuals.

To look for signs of potential positive selection acting on specific sites in the viral genome, we examined changes in the frequencies of recurring iSNVs over time. We plotted all detected instances of these shared iSNVs, even if they fell outside of the frequency range of 3% to 97% or fell below our chosen depth threshold of 1000 reads. Overall, the longitudinal dynamics of many iSNVs in both unvaccinated and immune individuals appeared highly stochastic, consistent with genetic drift and the absence of strong selection (**Figs 6,7; S2,S3**). Many iSNVs detected at high frequency at one or more timepoints fell below the limit of detection (LOD) at others within the same individual. In several of these cases, two or more iSNVs maintained highly similar frequencies over the course of infection, suggesting linkage (e.g. ORF1ab:V3261F and N:P67S in participant 451709; or UTR:t76a and UTR:t78g in participant 471588) (**Fig 6A,B**). The extreme fluctuations in frequency observed for some collection days may be explained in part by variation in the quality of population sampling associated with sample collection.

**Figure 6:**
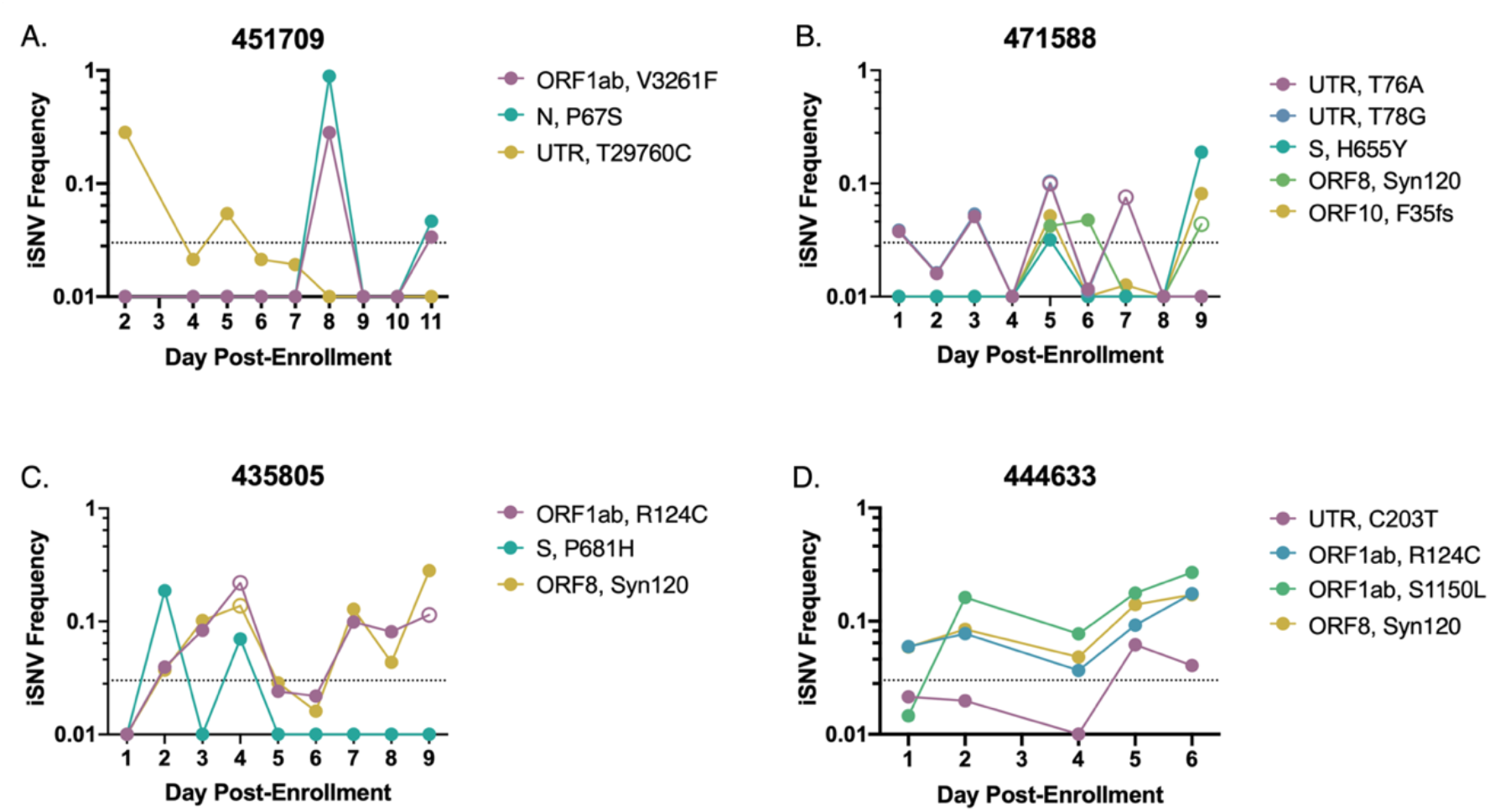
iSNV dynamics over time in saliva from naïve individuals. Frequency tracking of selected iSNVs from unvaccinated participants 451709 (A), 471588 (B), 435805 (C), and 444633 (D). Dashed line marks frequency threshold of 0.03. Unfilled points mark iSNVs with read depths below the threshold of 1000 reads.

**Figure 7:**
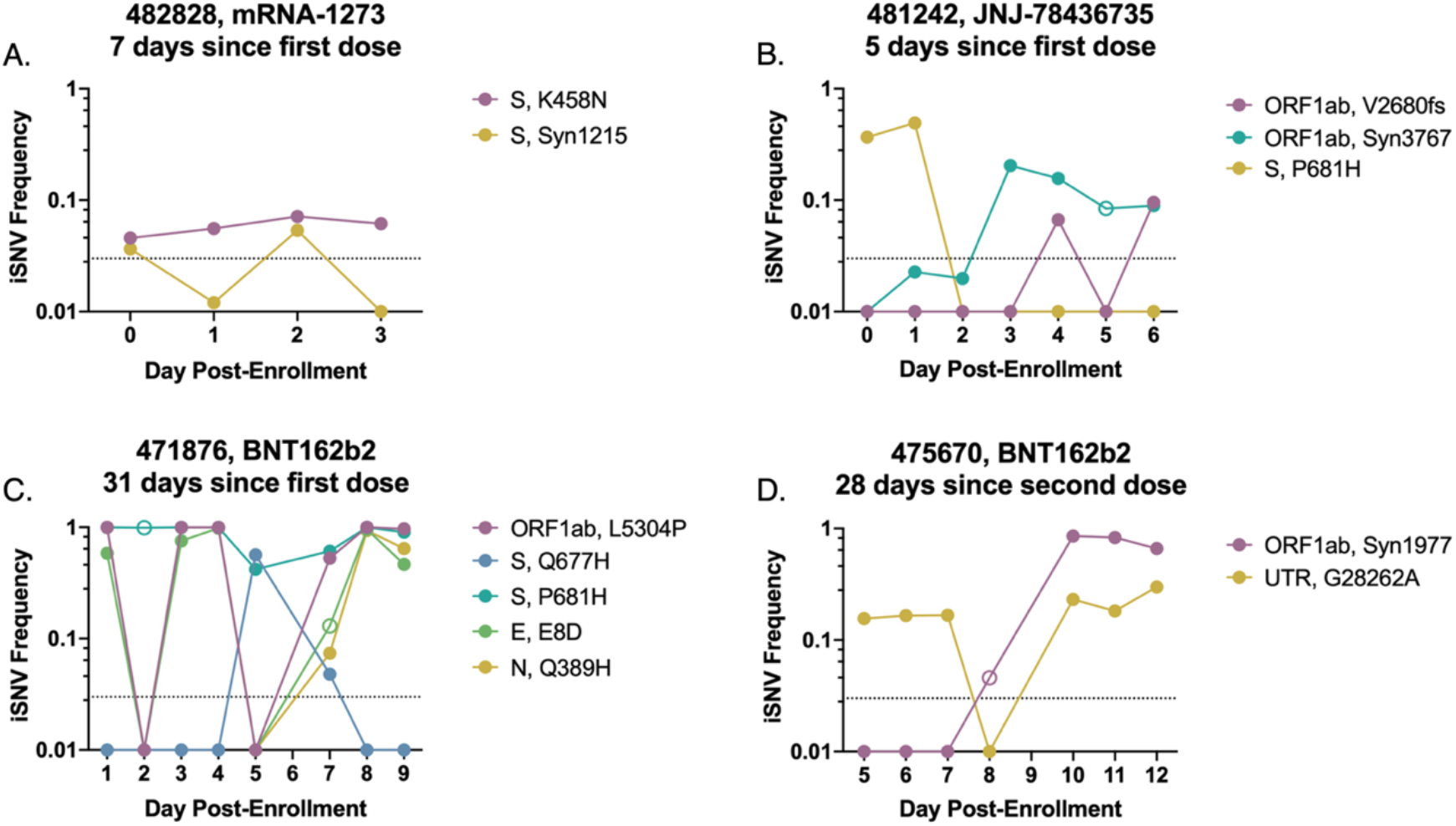
iSNV dynamics over time in saliva from vaccinated individuals. Frequency tracking of selected iSNVs from immune participants 482828 (newly vaccinated) (A), 481242 (newly vaccinated) (B), 471876 (partially vaccinated) (C), and 475670 (fully vaccinated) (D). Dashed line marks frequency threshold of 0.03. Unfilled points mark iSNVs with read depths below the threshold of 1000 reads. Panel headings indicate vaccine received and time between enrollment and last vaccine dose.

While most recurrent iSNVs did not appear to be under strong selection in naïve individuals, we observed several examples of variants that exhibited consistent patterns of emergence or decline over the course of acute infection that could be indicative of selection. We observed two ORF1ab substitutions (R124C and S1150L) in saliva that exhibited dynamics consistent with positive selection (**Fig 6C,D**). In the case of ORF1ab:R124C, these dynamics were observed across multiple individuals (**Fig S2**). Additionally, in nasal samples, we observed a substitution at ORF1ab:P5402H that emerged to near-fixation over the course of infection (**Fig S4**). However, across the global SARS-CoV-2 tree, these substitutions are observed either sporadically or not at all, suggesting the absence of positive selection at the between-host scale (**Fig S5**).

In participant 471588, the P.1. lineage-associated substitution S:H655Y showed up just above the LOD at day 5, dropped below the LOD, then re-emerged at over 10% on day 9 post-enrollment (**Fig 6B**). Another spike substitution associated with multiple variants of concern (Saxena et al., 2022, 2020a), S:P681H, was observed to emerge in participant 435805 on days 2 and 4, before ultimately falling back below the LOD. The within-host transience of a mutation associated with increased fitness at the global scale highlights how within-host evolutionary trends can diverge from global trends. Further illustrating the strength of genetic drift in these populations, we observed several cases in which synonymous mutations (which we assume to be neutral) rose to higher frequencies over the course of infection: *e.g*. the ORF1ab:1717 substitution in participant 435786 and the ORF1ab:1668 substitution in participant 451152 (**Fig S2**).

Overall, these data suggest that in a subset of acute infections, positive selection may be driving the emergence of specific iSNVs, but not generally to high enough frequencies to reliably transmit given the narrow transmission bottlenecks observed across multiple studies (Braun et al., 2021; Martin and Koelle, 2021; Valesano et al., 2021).

We did not observe any sweeps of antigenically significant spike substitutions in our small cohort of immune participants, suggesting that spike-based vaccination does not impose strong immune selection on the viral populations sampled over the course of acute infection (**Fig 7, S3**). The only recurrent, antigenically significant spike iSNV that we observed within our immune cohort resulted in a K458N substitution in the receptor binding domain (RBD), a residue that has been previously associated with monoclonal antibody (mAb) escape (**Fig 7A**) (Harvey et al., 2021; Liu et al., 2021b). This iSNV remained steady at a low frequency between 3% and 10% over the course of infection however, suggesting the absence of a strong selective advantage and low probability of forward transmission.

Outside of the established antigenic sites, we observed interesting dynamics near the S1/S2 cleavage site in immune individuals. In participant 481242, S:P681H was observed at middle frequencies on days 1 and 2 post enrollment but dropped below the LOD by day 3 and remained undetectable in later timepoints, suggesting a sweep by an S:P681 revertant (**Fig 7B**). In participant 471876, S:P681H was at or near fixation over the first four days of sampling, dropped in frequency on days 5 through 7, and then returned to near-fixation at day 8 (**Fig 7C**). On the two days where S:P681H dropped below 90%, a nearby spike substitution, Q677H, emerged to high frequency before dropping back below the LOD on days 8 and 9. The co-occurrence of the dip in S:P681H frequency with the emergence of S:Q677H and subsequent reversal are consistent with competition between these two substitutions. Both substitutions have proliferated at the global scale, suggesting that in some cases within-host and global dynamics may be aligned (Colson et al., 2022; Ghosh et al., 2021; Hodcroft et al., 2021; Saxena et al., 2022). Critically, S:Q677H peaked at a time (day 5 post-enrollment) when this participant was still viral culture positive (see figure 1 in (Ke et al., 2022b)), indicating the potential for this *de novo* variant to be successfully transmitted.

Overall, we did not detect obvious signs of antibody-mediated immune selection within immune individuals and found that iSNV frequencies often appeared to vacillate stochastically, like what was observed in naïve individuals. In participant 471876, several iSNVs (ORF1ab:L5304P, E:E8D, N:Q389H) fluctuated between fixation and frequencies below the LOD over the course of infection (**Fig 7C**). Additionally, in participant 475670, we observed a synonymous mutation (ORF1ab:Syn1977) rapidly rise to fixation (**Fig 7D**). However, it is hard to ascribe all observed dynamics entirely to genetic drift. Our observations of wild-type reversion and competition between iSNVs at the S1/S2 cleavage site (**Fig 7B,C**) suggest that selection may drive the within-host fluctuations of iSNVs at non-antigenic sites during acute infection of some immune individuals.

Finally, in keeping with our compartmentalization analysis, we found that frequencies of shared iSNVs in nasal swab samples over the course of infection often varied from the dynamics observed in saliva samples (**Fig S4**). Some of these differences arose from the lack of detection of a mutation in a certain environment, and some resulted when an iSNV that fluctuated in one environment was fixed in the other—for example, S:Y145del in participant 450241 was fixed in all nasal samples and was therefore only called as an iSNV in saliva samples (**Fig S2,4**). Following the trend observed in saliva samples, iSNV dynamics in nasal swab samples were generally stochastic, with only rare instances of dynamics consistent with selection.

## Discussion

By analyzing longitudinal samples collected daily over the course of acute infection, we captured a high-resolution temporal profile of SARS-CoV-2 within-host dynamics in humans. In general, we observed little evidence of strong selection acting on within-host viral populations in our cohort, consistent with previous reports (Braun et al., 2021; Tonkin-Hill et al., 2021; Valesano et al., 2021). This was true even within our group of vaccinated or previously infected individuals, mirroring a previous study of influenza virus that failed to detect the emergence of antigenic variants during infection of immune individuals (Debbink et al., 2017). These data suggest that respiratory viruses like SARS-CoV-2 and influenza virus may be able to replicate at some mucosal sites with minimal restriction by neutralizing antibodies, even within individuals with robust systemic antibody responses.

While signs of strong positive selection were rare in this cohort, we did identify a handful of nonsynonymous substitutions (S:Q677H, N:P67S, ORF1ab:P5402H) that emerged from below the limit of detection to high frequency over the course of infection. Importantly, S:Q677H emerged to 56.5% frequency on a day when the associated study participant had detectable infectious virus in a nasal swab (Ke et al., 2022b), suggesting the potential for this iSNV to be transmitted forward. Substitutions at S:Q677 (including Q677H) have independently emerged in multiple viral sub-lineages around the world, supporting that mutations at this site can be advantageous. We also observed signs of competition between S:Q677H and S:P681H within the same individual, with S:Q677H briefly emerging to a high frequency on a day at which the initially fixed S:P681H dipped in frequency. However, the observed reversion to a S:P681H-only genotype after day 7 suggests that the selective advantage conferred by S:P681H is greater than that of S:Q677H. This fitness advantage is supported globally by the more widespread proliferation of S:P681H-containing lineages in comparison to S:Q677H (Hadfield et al., 2018). Our data demonstrate how, in rare cases, *de novo* generated variants can emerge to within-host frequencies sufficient for transmission during acute infection.

In addition to the limited number of iSNVs that emerged to high frequency within single individuals, we observed several iSNVs that arose above background in multiple participants. Mapping the genomic locations of these shared mutations revealed several hotspots of non-synonymous mutation accumulation that differed between naïve and immune individuals. In naïve individuals, we identified hotspots at residues 402-457 in ORF1ab, and 655-681 in spike. The latter is especially interesting, as this region is directly adjacent to the S1/S2 cleavage site. Substitutions that modulate cleavage efficiency are important for transmission in ferrets (Peacock et al., 2021) and replication in cell culture (Johnson et al., 2021). S1/S2 cleavage site substitutions are characteristic features of the Omicron, Delta, and Alpha lineages, and have been shown to be responsible for Delta’s increased relative fitness compared with Alpha (Liu et al., 2021a), suggesting the importance of this domain in the adaptation of SARS-CoV-2 to humans. The enrichment of amino acid substitutions immediately upstream of the S1/S2 cleavage site in our cohort is further evidence that this region may be subject to stronger within-host selection in humans.

We also observed a surprisingly high density of N gene substitutions in immune participants. An observed hotspot of mutation accumulation in N:199-204 matches up with previous observations of frequent changes at positions 201-205 in the serine-rich region of the gene, several of which are characteristic of emerging lineages. R203K and G204R substitutions (both of which we observed in our samples) can increase the relative fitness of the virus, potentially through increased phosphorylation of the nucleocapsid (Johnson et al., 2022). These substitutions have also been associated with the transcription of an alternate subgenomic mRNA with anti-interferon activity (Mears et al., 2022). While spike protein substitutions are clearly the primary drivers of SARS-CoV-2 adaptation to humans, our results are also consistent with previous data suggesting an important role for the N gene during human adaptation.

Finally, several shared mutations occurred within untranslated regions of the viral genome. The most frequent of these was a t29760c substitution (in the 3’ UTR), which reoccurred across 9 different naïve individuals. We also observed recurring substitutions in the 5’ UTR, and, in immune individuals, a recurring substitution and insertion in the untranslated region preceding the N gene. The untranslated regions of the coronavirus genome form secondary structures that play a role in viral replication and translation (Yang and Leibowitz, 2015). It remains to be seen whether the recurring UTR mutations that we observe have appreciable effects on viral fitness.

Variant dynamics in multiple participants exhibited extreme fluctuations where iSNVs at or near fixation abruptly fall below the limit of detection, only to return to high frequencies days later. Given the abruptness of these fluctuations, it is doubtful that they were selection-driven. They could potentially be explained if there is a significant degree of spatial structuring of within-host viral genetic diversity, as has recently been described for influenza virus (Amato et al., 2022). Spatial structuring could promote more extreme, drift-driven fluctuations in sampled iSNV frequencies, due to bottleneck effects (Amato et al., 2022; Orton et al., 2020; Pfeiffer and Kirkegaard, 2006). Alternatively, these fluctuations might be artifactual, potentially arising from poor quality sampling of the viral population. We think the latter is unlikely, due to the Ct value thresholds we used for including samples in our analyses, but we cannot formally rule this possibility out. Regardless, either explanation further emphasizes the advantages of longitudinal sampling, as single-timepoint snapshots of viral populations can present misleading views of the within-host landscape.

Supporting the possibility that stochastic within-host SNV dynamics may partially result from spatial structuring, we observed significant compartmentalization between the oral and nasal environments over the course of SARS-CoV-2 infection in a subset of individuals. iSNVs varied in frequency between the two environments, a finding that builds on previous observations that peaks in viral shedding are often offset by several days between saliva and nasal environments (Ke et al., 2022a), and that shedding is sometimes limited to the saliva compartment in immune individuals (Ke et al., 2022b). These results suggest the potential for tissue-specific adaptation by the virus and reaffirm that sampling of a single tissue site may not provide a complete view of viral population diversity within a host.

A clear advantage of repeated longitudinal sampling is that it allows for higher confidence variant calling compared with single-timepoint sampling. Across individual samples, we measured iSNV counts ranging from zero to several hundred and found that these values could shift rapidly within an individual over short periods of time. However, the number of variants shared across multiple days was consistent across both cohorts and remained relatively low, with both groups exhibiting shared iSNV counts that align with previous assessments of within-host diversity (Valesano et al. 2021; Lythgoe et al. 2021).

Altogether, our results suggest that viral evolution is largely driven by stochastic forces during acute infections but that in rare occasions selection can drive the emergence of iSNVs capable of forward transmission. Furthermore, our recurrent detection of iSNVs that have been successful (or not) at the global scale indicate areas of alignment and discordance between within-host and between-host selective pressures and thus help shed light on the forces that shape global patterns of SARS-CoV-2 evolution.

## Acknowledgments

This work has been generously supported by the National Heart, Lung, and Blood Institute of the National Institutes of Health award 3U54HL143541-02S2) through the RADx-Tech program to D.D.M., L.L.G. and C.B.B., as well as additional funds generously provided by the Carl R. Woese Institute for Genomic Biology and the Department of Microbiology at the University of Illinois at Urbana-Champaign.

We are grateful to Alvaro Hernandez, Chris Wright, and the DNA services team at the Roy J. Carver Biotechnology Center for assistance in next generation sequencing.

We gratefully acknowledge the Authors from the originating laboratories responsible for obtaining the specimens and the submitting laboratories where genetic sequence data were generated and shared via the GISAID Initiative, on which Fig S5 is based (**Table S1**)

## Declaration of interests

The authors declare no competing interests.

## Methods

### Sample collection

To monitor on-campus COVID cases, students and employees at the University of Illinois at Urbana-Champaign were required to submit biweekly saliva samples for RTqPCR testing. Individuals who tested positive were given the option to enroll in a longitudinal sample collection study within 1 day of receiving a positive result. Additionally, individuals who been in close contact with a positive case were eligible to enroll in the same study within 5 days of their exposure. Enrolled participants then provided saliva samples and mid-turbinate nasal swabs for 14 days after the date of their first positive test (This collection protocol is described in detail in (Ke et al., 2022a)) Within 12 hours of collection, RTqPCR was performed on heat-inactivated saliva samples to assess viral load, as described in Ranoa et al. (Ranoa et al., 2020). Nasal swab samples were stored in viral transport media at −80° C and shipped to Johns Hopkins University for analysis.

Participants were designated as fully vaccinated if they had been infected at least 14 days after receiving a single-dose vaccine (JNJ-78436735) or a second dose of a two-dose vaccine (BNT162b2 or mRNA-1273). If at least 14 days had passed since receiving the first dose of a two-dose vaccine, participants were designated as partially vaccinated, and if less than 14 days had passed since receiving a dose of any vaccine, participants were designated as newly vaccinated. Study enrollment was concluded prior to the approval of vaccine boosters.

### Participant selection

After RTqPCR analysis, unvaccinated participants with fewer than three saliva samples under the cycle threshold (Ct) cutoff value of 28 were filtered from the dataset. The remaining participants were sorted and ranked based on their number of quality samples and the range of dates covered by these samples. The top 20 participants were selected for further analysis. All immune participants with saliva samples under a Ct value of 30 were retained, which resulted in a study group of 12 individuals. From this combined cohort of naïve and immune individuals, nasal samples from 14 individuals were chosen to evaluate environmental differences between the oral and nasal cavities.

### RNA extraction and sequencing (Saliva samples)

To extract viral RNA, a volume of 140 uL from each heat-inactivated saliva sample was processed using the QIAamp viral RNA mini kit. Viral cDNA was generated from 100 ng of the resulting RNA aliquots and sequencing libraries were prepared from the cDNA using the Swift SNAP Amplicon SARS-CoV-2 kit. Deep sequencing was then performed on an Illumina NovaSeq. Raw sequences were processed using the nf-core/viral-recon workflow, in order to align sequences to the Wuhan-Hu-1 reference genome and extract frequencies and annotations for variants at frequencies higher than 0.01. Lineages were assigned using *Pango* version 1.2.34.

### Analysis of variant dynamics (Saliva samples)

Variants were extracted from sequences aligned to Wuhan-Hu-1 using *iVar,* and variant effects were annotated with *SnpEff*. To focus on minor sequence variants, variants present above a frequency of 0.97 were left out of the dataset. Variants at frequencies lower than 0.03 were also removed to decrease the potential for error due to noise. A per-nucleotide depth threshold of 1000 reads was also applied to the dataset. For each participant, variants present across two or more days of infection were extracted, and their frequencies tracked. Though the depth and frequency cutoffs described above were used to identify these shared variants, frequency tracking was performed on a dataset curated without thresholds, to avoid cases in which variants crossing either threshold may erroneously appear to fall out of the dataset. Variants with per-nucleotide coverage values below the cutoff were specially marked and plotted to indicate their low depth values. *SnpEff* annotations were used to characterize shared variants as synonymous, nonsynonymous, or untranslated, and to assign them to the appropriate region of the SARS-CoV-2 genome. Genome positions of variants were visualized using *trackViewer* package (version 1.28.1) in R (Ou and Zhu, 2019). The ratio of nonsynonymous to synonymous variants (dN/dS) was then calculated for each sample, using all variants above the depth threshold of 1000. Variant counts were normalized to an estimate of the number of nonsynonymous sites (9803) or synonymous sites (19606) in the genome, which were calculated by estimating that each codon contains one synonymous site. Infinite and NaN values were excluded from further analysis.

### RNA extraction and sequencing (Nasal swab samples)

Mid-turbinate nasal swab specimen aliquots were maintained at −80°C prior to use. RNA was extracted from 300 μl of clinical specimen using the Chemagic™ 360 system (Perkin Elmer) according to the manufacturer’s specifications. RNA was eluted with 60μl elution buffer and stored at −80°C until use. cDNA synthesis was performed using Superscript IV reverse transcriptase Kit (ThermoFisher Scientific) following the manufacturer’s protocol. The amplification of the genome was performed using Q5 Hot Start DNA Polymerase (ThermoFisher) and two pools of primers, each containing unique non-overlapping binding sites covering half of the SARS-CoV-2 genome. Library preparation was performed following the protocol provided with the Illumina Nextera DNA Flex kit for sample inputs of 100-500ng.

Briefly, adapter sequences were ligated to genomic DNA fragments via Tagmentation, after which mean amplicon size was determined via Agilent TapeStation 4500 and concentration was determined using a Qubit Flex Fluorometer (Invitrogen). Size normalization was performed and samples were diluted to a loading concentration of 1.2-1.3pM. Samples were sequenced using Illumina MiniSeq High Output Reagents (150 cycles).

### Analysis of genetic compartmentalization between sample sites

To account for sequencing differences between saliva samples and nasal swab samples (which were processed at different facilities), we imposed varying quality control thresholds on each dataset. To remove potentially artefactual iSNVs, we imposed a per-nucleotide depth cutoff of 500 reads in saliva samples and 200 reads in nasal swab samples. To remove samples with low overall coverage, we imposed a mean coverage cutoff of 1000 on saliva samples and a median coverage cutoff of 200 on nasal samples. We assigned SNPs present below a threshold of 1% a frequency of 0, and those present above a threshold of 99% a frequency of 1. Because nasal swab samples were sequenced using ARTIC primers, we also filtered out common SNP artifacts that arise frequently with this primer set (2020b). In one individual (451709), we also excluded samples from days 4 and 7, due to evidence of possible cross-contamination. For each participant, pairwise F_ST_ values were calculated for all possible pairs of samples, including pairs of saliva samples, pairs of nasal samples, and pairs of one saliva sample and one nasal sample.

### Phylogenetic analysis

The metadata file for all sequences present in the GISAID EpiCov database (10.2807/1560-7917.ES.2017.22.13.30494) was downloaded on June 10th, 2022. This metadata file was was filtered to include only entries from human hosts, only complete and high coverage entries, and only those with complete sampling dates. The filtered metadata entries were downsampled to at most 100 per month. Downsampling was conducted in Python v3.9.4 (Van Rossum and Drake, 2009)using Pandas v1.1.4 (10.5281/zenodo.3509134) and Numpy v1.19.4 (10.1038/s41586-020-2649-2). The selected sequences were downloaded from GISAID EpiCov and aligned to Wuhan/WIV04 (EPI_ISL_402124) using MAFFT v7.464 (10.1093/molbev/mst010), removing any insertions to the reference. IQtree v2.1.3 (10.1093/molbev/msaa015) was used to infer a phylogenetic tree of the aligned sequences using a GTR+G4 substitution model, saving with Wuhan/WIV04 as the outgroup. TreeTime v0.8.0 (10.1093/ve/vex042) was used to filter sequences to include only those falling within four interquartile ranges of the best fit molecular clock, rooting at Wuhan/WIV04. Wuhan/WIV04 was forced to be included in the filtered tree and no other tips were identified as failing this filter.

For each of the amino acids of interest (ORF1ab R124, ORF1ab S1150, ORF1ab P5402), we first identified the corresponding nucleotide positions and then identified the nucleotide identity at each of those sites for each sequence in the alignment. These nucleotide identities were used to infer the amino acid for each sequence at each position. Note that this method does not account for the presence of frame shift mutations, however, we expect these to be sufficiently rare as to not bias our results.

For each of the four substitutions we plotted the downsampled phylogenetic tree, labeling any tips with amino acids that did not match the reference. Any tips in which any of the nucleotides in the codon of interest were deleted or ambiguously genotyped were ignored. Visualization was done in Python using Matplotlib v3.5.1 (10.1109/MCSE.2007.55) and Baltic v0.1.5 [https://github.com/evogytis/baltic].

### Substitution frequency analysis

The GISAID EpiCoV “MSA full” alignment was downloaded on June 2nd, 2022. All sequences in this file have been aligned to Wuhan/WIV04 using MAFFT, retaining any insertions relative to the reference. Full details on how this file were generated are available from GISAID.

For each of the four amino acid positions of interest we first identified the corresponding nucleotide positions in the gapped alignment and identified the nucleotides at each of these for each position. These nucleotides were used to infer the amino acid for each sequence in the full alignment at each position. Similar to above, this method does not account for frameshift mutations.

For each amino acid position, we identified the percentage of sequences harboring non-reference amino acids per month, ignoring any sequences in each one of the nucleotides was deleted or ambiguously genotyped. All amino acid identifies with a maximum monthly frequency less than or equal to 0.01% were grouped into an “Other” category. This analysis was conducted in Python using Pandas and visualized with Matplotlib.

## Supplemental Figures

**Supplemental Figure 1:**
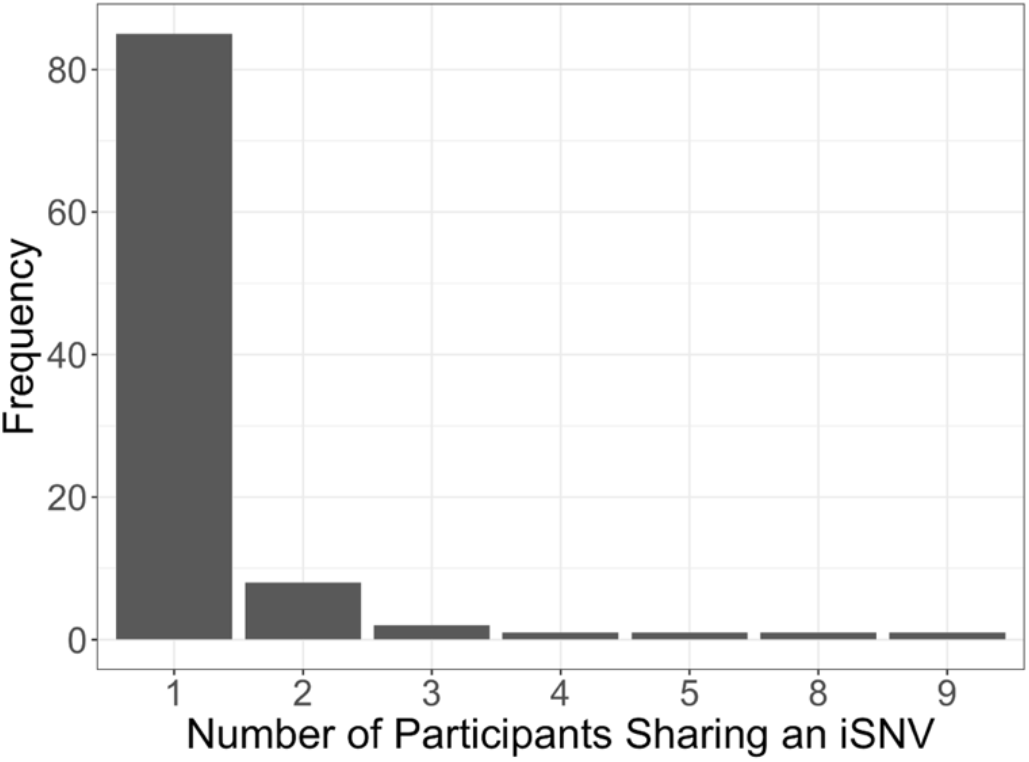
Distribution of shared iSNVs found across multiple naive participants. No shared iSNVs were detected in more than 2 vaccinated participants.

**Supplemental Figure 2:**
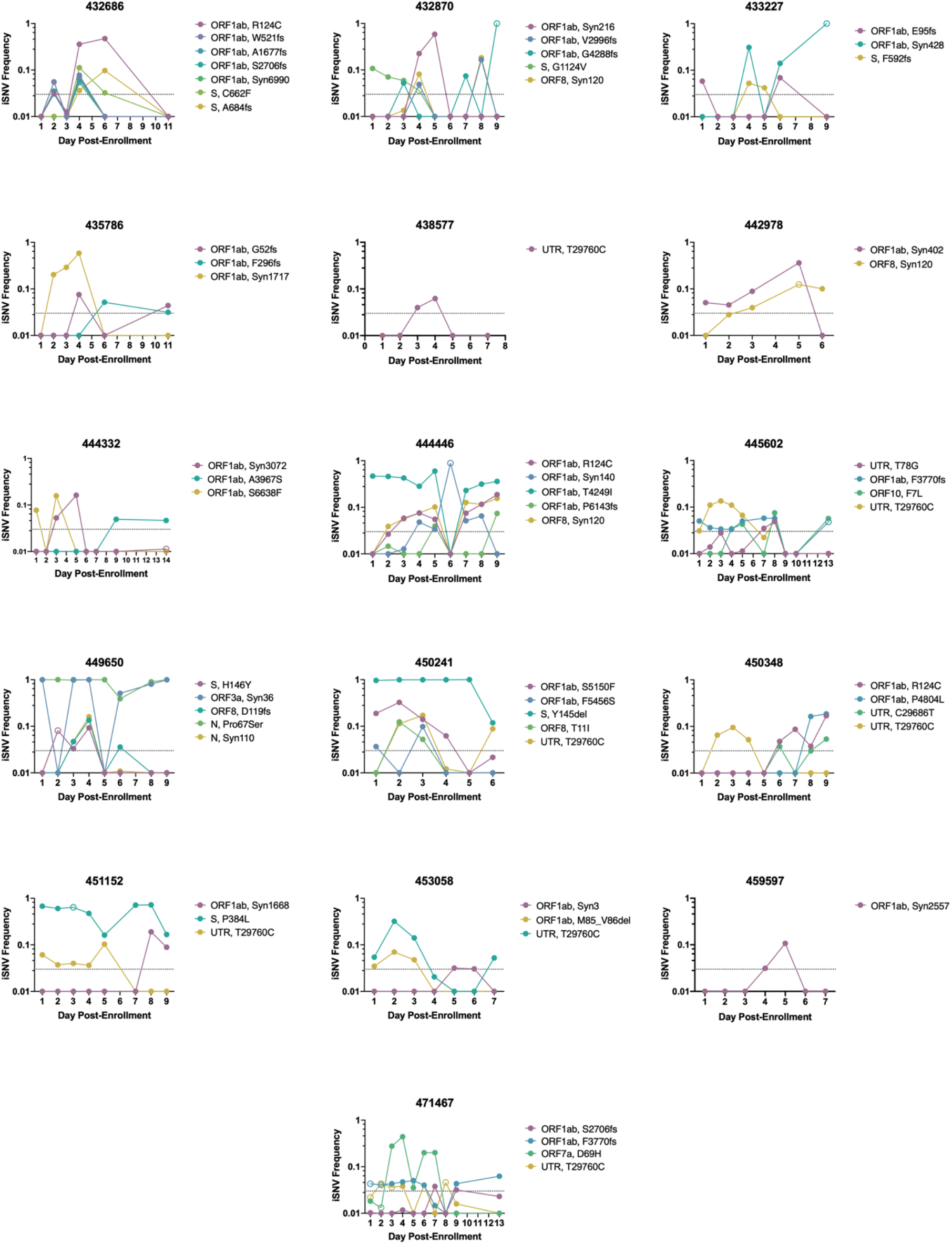
iSNV dynamics over time from all unvaccinated participants. Frequency tracking of selected iSNVs in unvaccinated participants. Dashed line marks frequency threshold of 0.03. Unfilled points mark iSNVs with read depths below the threshold of 1000 reads.

**Supplemental Figure 3:**
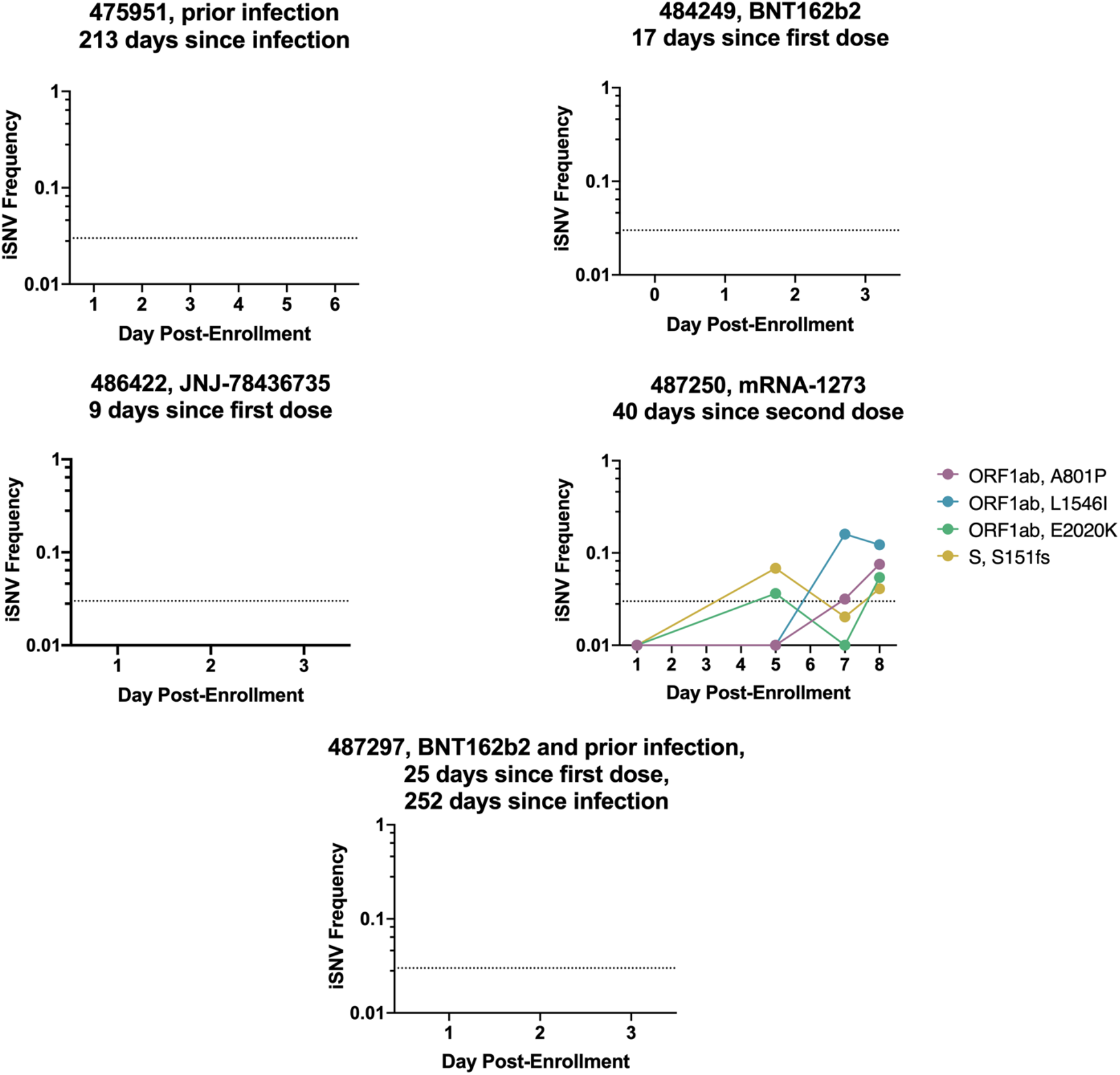
iSNV dynamics over time from all immune participants. Frequency tracking of selected iSNVs in immune participants. Dashed line marks frequency threshold of 0.03. Unfilled points mark iSNVs with read depths below the threshold of 1000 reads.

**Supplemental Figure 4:**
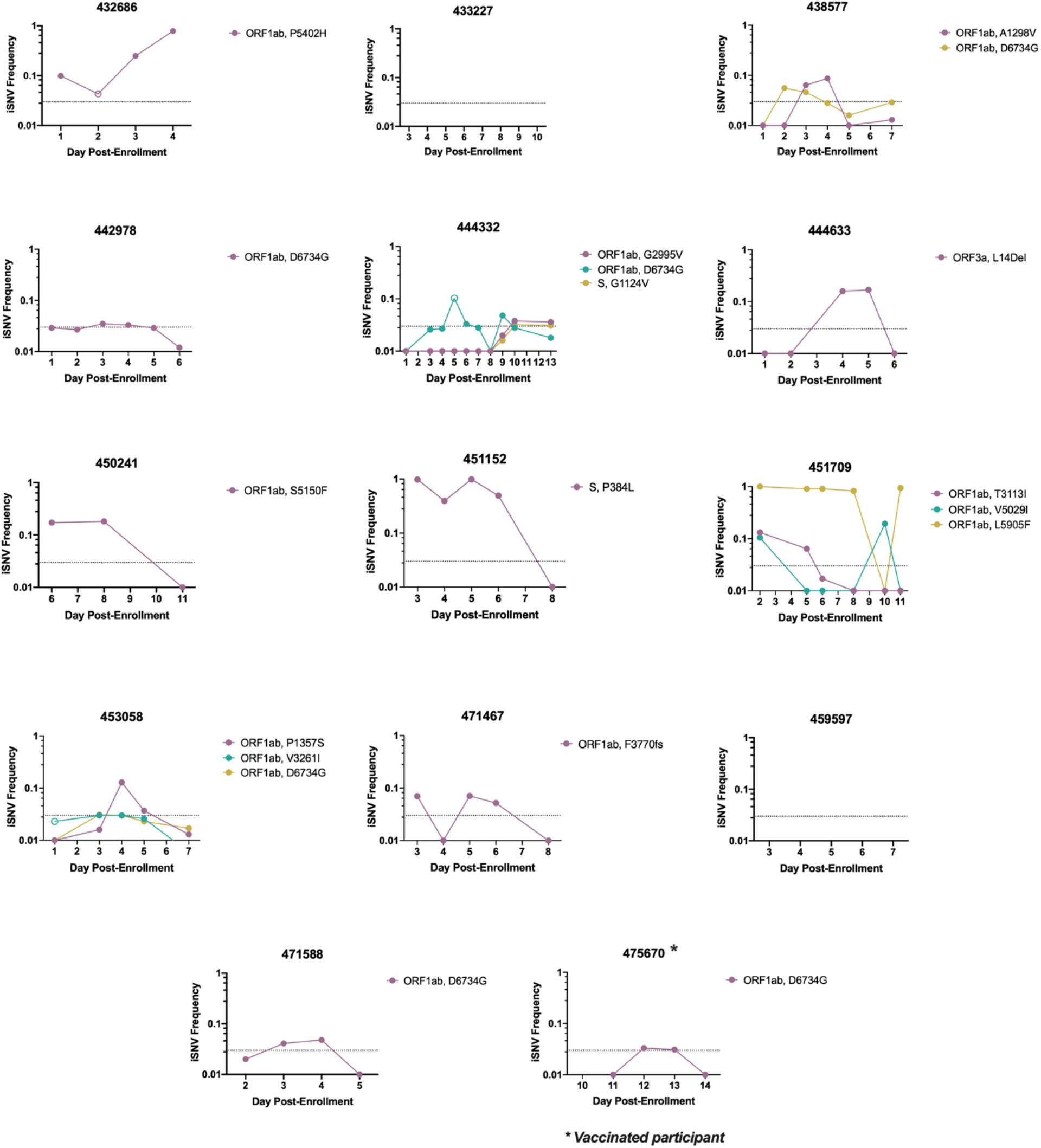
iSNV dynamics over time from all nasal samples. Frequency tracking of selected iSNVs from nasal swab samples. Dashed line marks frequency threshold of 0.03. Unfilled points mark iSNVs with read depths below the threshold of 1000 reads.

**Supplemental Figure 5.**
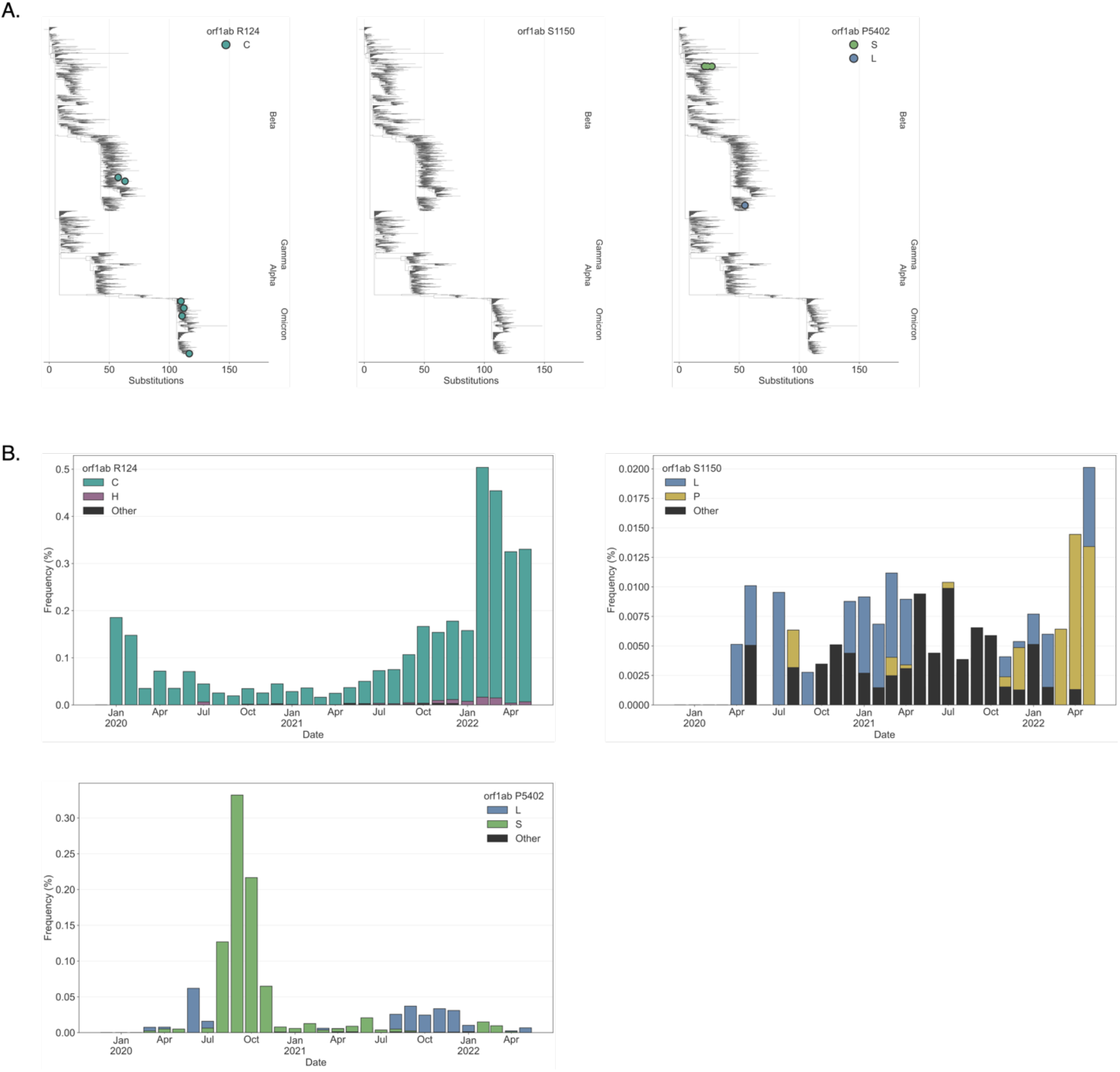
Global occurrence of selected iSNVs. **(A)** iSNV-associated amino acid changes plotted on a downsampled (100 sequences per month) global phylogeny of SARS-CoV-2 sequences. **(B)** Frequency of iSNV-associated amino acid changes from January 2020 to April 2022 in all quality filtered SARS-CoV-2 sequences.

## Notes

### Competing Interest Statement

The authors have declared no competing interest.

### Summary of Updates

Added supplemental table with acknowledgment of sources of GISAID sequence data used. Corrected some references, typos, and added a missing author affiliation.

## References

Amato, K.A., Haddock, L.A., Braun, K.M., Meliopoulos, V., Livingston, B., Honce, R., Schaack, G.A., Boehm, E., Higgins, C.A., Barry, G.L., et al. (2022). Influenza A virus undergoes compartmentalized replication in vivo dominated by stochastic bottlenecks. Nat Commun 13, 3416. https://doi.org/10.1038/s41467-022-31147-0.

Avanzato, V.A., Matson, M.J., Seifert, S.N., Pryce, R., Williamson, B.N., Anzick, S.L., Barbian, K., Judson, S.D., Fischer, E.R., Martens, C., et al. (2020). Case Study: Prolonged Infectious SARS-CoV-2 Shedding from an Asymptomatic Immunocompromised Individual with Cancer. Cell 183, 1901–1912.e9. https://doi.org/10.1016/j.cell.2020.10.049.

Baang, J.H., Smith, C., Mirabelli, C., Valesano, A.L., Manthei, D.M., Bachman, M.A., Wobus, C.E., Adams, M., Washer, L., Martin, E.T., et al. (2021). Prolonged Severe Acute Respiratory Syndrome Coronavirus 2 Replication in an Immunocompromised Patient. The Journal of Infectious Diseases 223, 23–27. https://doi.org/10.1093/infdis/jiaa666.

Braun, K.M., Moreno, G.K., Wagner, C., Accola, M.A., Rehrauer, W.M., Baker, D.A., Koelle, K., O’Connor, D.H., Bedford, T., Friedrich, T.C., et al. (2021). Acute SARS-CoV-2 infections harbor limited within-host diversity and transmit via tight transmission bottlenecks. PLOS Pathogens 17, e1009849. https://doi.org/10.1371/journal.ppat.1009849.

Choi, B., Choudhary, M.C., Regan, J., Sparks, J.A., Padera, R.F., Qiu, X., Solomon, I.H., Kuo, H.-H., Boucau, J., Bowman, K., et al. (2020). Persistence and Evolution of SARS-CoV-2 in an Immunocompromised Host. N Engl J Med 383, 2291–2293. https://doi.org/10.1056/NEJMc2031364.

Cobey, S., Larremore, D.B., Grad, Y.H., and Lipsitch, M. (2021). Concerns about SARS-CoV-2 evolution should not hold back efforts to expand vaccination. Nature Reviews Immunology 1–6. https://doi.org/10.1038/s41577-021-00544-9.

Colson, P., Delerce, J., Burel, E., Beye, M., Fournier, P.-E., Levasseur, A., Lagier, J.-C., and Raoult, D. (2022). Occurrence of a substitution or deletion of SARS-CoV-2 spike amino acid 677 in various lineages in Marseille, France. Virus Genes 58, 53–58. https://doi.org/10.1007/s11262-021-01877-2.

Corey, L., Beyrer, C., Cohen, M.S., Michael, N.L., Bedford, T., and Rolland, M. (2021). SARS-CoV-2 Variants in Patients with Immunosuppression. New England Journal of Medicine 385, 562–566. https://doi.org/10.1056/NEJMsb2104756.

Debbink, K., McCrone, J.T., Petrie, J.G., Truscon, R., Johnson, E., Mantlo, E.K., Monto, A.S., and Lauring, A.S. (2017). Vaccination has minimal impact on the intrahost diversity of H3N2 influenza viruses. PLOS Pathogens 13, e1006194. https://doi.org/10.1371/journal.ppat.1006194.

Ghosh, A.K., Kaiser, M., Molla, Md.M.A., Nafisa, T., Yeasmin, M., Ratul, R.H., Sharif, Md.M., Akram, A., Hosen, N., Mamunur, R., et al. (2021). Molecular and Serological Characterization of the SARS-CoV-2 Delta Variant in Bangladesh in 2021. Viruses 13, 2310. https://doi.org/10.3390/v13112310.

Hadfield, J., Megill, C., Bell, S.M., Huddleston, J., Potter, B., Callender, C., Sagulenko, P., Bedford, T., and Neher, R.A. (2018). Nextstrain: real-time tracking of pathogen evolution. Bioinformatics 34, 4121–4123. https://doi.org/10.1093/bioinformatics/bty407.

Harvey, W.T., Carabelli, A.M., Jackson, B., Gupta, R.K., Thomson, E.C., Harrison, E.M., Ludden, C., Reeve, R., Rambaut, A., Peacock, S.J., et al. (2021). SARS-CoV-2 variants, spike mutations and immune escape. Nat Rev Microbiol 19, 409–424. https://doi.org/10.1038/s41579-021-00573-0.

Hodcroft, E.B., Domman, D.B., Snyder, D.J., Oguntuyo, K.Y., Diest, M.V., Densmore, K.H., Schwalm, K.C., Femling, J., Carroll, J.L., Scott, R.S., et al. (2021). Emergence in late 2020 of multiple lineages of SARS-CoV-2 Spike protein variants affecting amino acid position 677. 2021.02.12.21251658. https://doi.org/10.1101/2021.02.12.21251658.

Johnson, B.A., Xie, X., Bailey, A.L., Kalveram, B., Lokugamage, K.G., Muruato, A., Zou, J., Zhang, X., Juelich, T., Smith, J.K., et al. (2021). Loss of furin cleavage site attenuates SARS-CoV-2 pathogenesis. Nature 591, 293–299. https://doi.org/10.1038/s41586-021-03237-4.

Johnson, B.A., Zhou, Y., Lokugamage, K.G., Vu, M.N., Bopp, N., Crocquet-Valdes, P.A., Kalveram, B., Schindewolf, C., Liu, Y., Scharton, D., et al. (2022). Nucleocapsid mutations in SARS-CoV-2 augment replication and pathogenesis. PLoS Pathog 18, e1010627. https://doi.org/10.1371/journal.ppat.1010627.

Ke, R., Martinez, P.P., Smith, R.L., Gibson, L.L., Mirza, A., Conte, M., Gallagher, N., Luo, C.H., Jarrett, J., Zhou, R., et al. (2022a). Daily longitudinal sampling of SARS-CoV-2 infection reveals substantial heterogeneity in infectiousness. Nat Microbiol 7, 640–652. https://doi.org/10.1038/s41564-022-01105-z.

Ke, R., Martinez, P.P., Smith, R.L., Gibson, L.L., Achenbach, C.J., McFall, S., Qi, C., Jacob, J., Dembele, E., Bundy, C., et al. (2022b). Longitudinal analysis of SARS-CoV-2 vaccine breakthrough infections reveal limited infectious virus shedding and restricted tissue distribution. Open Forum Infectious Diseases ofac192. https://doi.org/10.1093/ofid/ofac192.

Kemp, S.A., Collier, D.A., Datir, R.P., Ferreira, I.A.T.M., Gayed, S., Jahun, A., Hosmillo, M., Rees-Spear, C., Mlcochova, P., Lumb, I.U., et al. (2021). SARS-CoV-2 evolution during treatment of chronic infection. Nature 592, 277–282. https://doi.org/10.1038/s41586-021-03291-y.

Kryazhimskiy, S., and Plotkin, J.B. (2008). The Population Genetics of dN/dS. PLoS Genet 4, e1000304. https://doi.org/10.1371/journal.pgen.1000304.

Liu, Y., Liu, J., Johnson, B.A., Xia, H., Ku, Z., Schindewolf, C., Widen, S.G., An, Z., Weaver, S.C., Menachery, V.D., et al. (2021a). Delta spike P681R mutation enhances SARS-CoV-2 fitness over Alpha variant.

Liu, Z., VanBlargan, L.A., Bloyet, L.-M., Rothlauf, P.W., Chen, R.E., Stumpf, S., Zhao, H., Errico, J.M., Theel, E.S., Liebeskind, M.J., et al. (2021b). Identification of SARS-CoV-2 spike mutations that attenuate monoclonal and serum antibody neutralization. Cell Host & Microbe 29, 477–488.e4. https://doi.org/10.1016/j.chom.2021.01.014.

Lythgoe, K.A., Hall, M., Ferretti, L., de Cesare, M., MacIntyre-Cockett, G., Trebes, A., Andersson, M., Otecko, N., Wise, E.L., Moore, N., et al. (2021). SARS-CoV-2 within-host diversity and transmission. Science 372, eabg0821. https://doi.org/10.1126/science.abg0821.

Martin, M.A., and Koelle, K. (2021). Comment on “Genomic epidemiology of superspreading events in Austria reveals mutational dynamics and transmission properties of SARS-CoV-2.” Sci. Transl. Med. 13, eabh1803. https://doi.org/10.1126/scitranslmed.abh1803.

Mears, H.V., Young, G.R., Sanderson, T., Harvey, R., Crawford, M., Snell, D.M., Fowler, A.S., Hussain, S., Nicod, J., Peacock, T.P., et al. (2022). Emergence of new subgenomic mRNAs in SARS-CoV-2. 19..

Orton, R.J., Wright, C.F., King, D.P., and Haydon, D.T. (2020). Estimating viral bottleneck sizes for FMDV transmission within and between hosts and implications for the rate of viral evolution. Interface Focus 10, 20190066. https://doi.org/10.1098/rsfs.2019.0066.

Ou, J., and Zhu, L.J. (2019). trackViewer: a Bioconductor package for interactive and integrative visualization of multi-omics data. Nat Methods 16, 453–454. https://doi.org/10.1038/s41592-019-0430-y.

Peacock, T.P., Goldhill, D.H., Zhou, J., Baillon, L., Frise, R., Swann, O.C., Kugathasan, R., Penn, R., Brown, J.C., Sanchez-David, R.Y., et al. (2021). The furin cleavage site in the SARS-CoV-2 spike protein is required for transmission in ferrets. Nat Microbiol 6, 899–909. https://doi.org/10.1038/s41564-021-00908-w.

Pfeiffer, J.K., and Kirkegaard, K. (2006). Bottleneck-mediated quasispecies restriction during spread of an RNA virus from inoculation site to brain. Proceedings of the National Academy of Sciences 103, 5520–5525. https://doi.org/10.1073/pnas.0600834103.

Ranoa, D.R.E., Holland, R.L., Alnaji, F.G., Green, K.J., Wang, L., Brooke, C.B., Burke, M.D., Fan, T.M., and Hergenrother, P.J. (2020). Saliva-Based Molecular Testing for SARS-CoV-2 that Bypasses RNA Extraction. 2020.06.18.159434. https://doi.org/10.1101/2020.06.18.159434.

Ranoa, D.R.E., Holland, R.L., Alnaji, F.G., Green, K.J., Wang, L., Fredrickson, R.L., Wang, T., Wong, G.N., Uelmen, J., Maslov, S., et al. (2021). Mitigation of SARS-CoV-2 Transmission at a Large Public University (Infectious Diseases (except HIV/AIDS)).

Saad-Roy, C.M., Morris, S.E., Metcalf, C.J.E., Mina, M.J., Baker, R.E., Farrar, J., Holmes, E.C., Pybus, O.G., Graham, A.L., Levin, S.A., et al. (2021). Epidemiological and evolutionary considerations of SARS-CoV-2 vaccine dosing regimes. Science 372, 363–370. https://doi.org/10.1126/science.abg8663.

Saxena, S.K., Kumar, S., Ansari, S., Paweska, J.T., Maurya, V.K., Tripathi, A.K., and Abdel-Moneim, A.S. (2022). Characterization of the novel SARS-CoV-2 Omicron (B.1.1.529) variant of concern and its global perspective. Journal of Medical Virology 94, 1738–1744. https://doi.org/10.1002/jmv.27524.

Smith, R.L., Gibson, L.L., Martinez, P.P., Ke, R., Mirza, A., Conte, M., Gallagher, N., Conte, A., Wang, L., Fredrickson, R., et al. (2021). Longitudinal Assessment of Diagnostic Test Performance Over the Course of Acute SARS-CoV-2 Infection. The Journal of Infectious Diseases 224, 976–982. https://doi.org/10.1093/infdis/jiab337.

Tonkin-Hill, G., Martincorena, I., Amato, R., Lawson, A.R., Gerstung, M., Johnston, I., Jackson, D.K., Park, N., Lensing, S.V., Quail, M.A., et al. (2021). Patterns of within-host genetic diversity in SARS-CoV-2. ELife 10, e66857. https://doi.org/10.7554/eLife.66857.

Truong, T.T., Ryutov, A., Pandey, U., Yee, R., Goldberg, L., Bhojwani, D., Aguayo-Hiraldo, P., Pinsky, B.A., Pekosz, A., Shen, L., et al. (2021). Increased viral variants in children and young adults with impaired humoral immunity and persistent SARS-CoV-2 infection: A consecutive case series. EBioMedicine 67, 103355. https://doi.org/10.1016/j.ebiom.2021.103355.

Valesano, A.L., Rumfelt, K.E., Dimcheff, D.E., Blair, C.N., Fitzsimmons, W.J., Petrie, J.G., Martin, E.T., and Lauring, A.S. (2021). Temporal dynamics of SARS-CoV-2 mutation accumulation within and across infected hosts. PLOS Pathogens 17, e1009499. https://doi.org/10.1371/journal.ppat.1009499.

Van Rossum, G., and Drake, F.L. (2009). Python 3 Reference Manual (Scotts Valley, CA: CreateSpace).

Yang, D., and Leibowitz, J.L. (2015). The structure and functions of coronavirus genomic 3’ and 5’ ends. Virus Research 206, 120–133. https://doi.org/10.1016/j.virusres.2015.02.025.

(2020a). Preliminary genomic characterisation of an emergent SARS-CoV-2 lineage in the UK defined by a novel set of spike mutations - SARS-CoV-2 coronavirus / nCoV-2019 Genomic Epidemiology.

(2020b). Issues with SARS-CoV-2 sequencing data - SARS-CoV-2 coronavirus / nCoV-2019 Genomic Epidemiology.

