## Supplemental Table 1 for "Within-host evolutionary dynamics and tissue compartmentalization during acute SARS-CoV-2 infection"

**Data Availability**

GISAID Identifier: EPI_SET_20220621ms
DOI: https://doi.org/10.55876/gis8.220621ms

All genome sequences and associated metadata in this dataset are published in GISAID's EpiCoV database. To view the contributors of each individual sequence with details such as accession number, Virus name, Collection date, Originating Lab and Submitting Lab and the list of Authors, visit https://doi.org/10.55876/gis8.220621ms

**Data Snapshot**

- EPI_SET_20220621ms is composed of 5,285,238 individual genome sequences.
- The collection dates range from 2019-12-24 to 2022-06-04;
- Data were collected in 203 countries and territories;
- All sequences in this dataset are compared relative to hCoV-19/Wuhan/WIV04/2019 (WIV04), the official reference sequence employed by GISAID (EPI_ISL_402124). Learn more at https://gisaid.org/WIV04.
